# Mechanisms underlying the therapeutic effects of Semen Cuscutae in treating Recurrent Spontaneous Abortion based on Network Pharmacology and Molecular Docking

**DOI:** 10.1101/2023.05.28.542653

**Authors:** Wenfei Zheng, Manshu Lei, Yao Yao, Jingqiong Zhan, Yiming Zhang, Feng Huang

**Affiliations:** Department of Gynecology and Obstetrics, the People’s Hospital of China Three Gorges University/the First People’s Hospital of Yichang, Yichang 443000, China

**Author notes:** Jiefang Road, NO 2, Yichang, Hubei province, China. Correspondence (WZ). These authors contributed equally to this work (Wenfei Zheng, Manshu Lei).

**Keywords:** immune-inflammatory, *Semen Cuscutae*, recurrent spontaneous abortion, network pharmacology, molecular docking, tumor necrosis factor-alpha inhibitor

## Abstract

**Background:** This paper aims to analyze the active components of SC by network pharmacology and screen the most stable compounds with TNF-a by molecular docking, to explore the mechanism of SC treatment of RSA and provide theoretical basis for drug development.

**Methods:** Active compounds of *SC* and the potential inflammatory targets of RSA were obtained from the Traditional Chinese Medicine Systematic Pharmacology Database and GeneCard, respectively. The interaction network of RSA–*SC*-target gene was accomplished and Visualizing by STRING database and Cytoscape software.GO and KEGG pathway enrichment analyses were obtained from DAVID to further explore the RSA mechanism and therapeutic effects of *SC*. Interactions between TNF-α and active compounds were analyzed by molecular docking.

**Results:** 10 active ingredients and 128 target genes were screened from *SC*, and 79 of them intersected with RSA target inflammatory genes,which were considered to be potential therapeutic targets. Network pharmacological analysis showed that Sesamin, matrine, matrol and other SC active components had good correlation with the inflammatory target genes of RSA.Related genes include PGR, PTGS1, PTGS2, TGFB1 and CHRNA7. Multiple signaling pathways are involved in RSA pathogenesis,sunh as TNF-α signaling pathway, HIF-1 signaling pathway, estrogen signaling pathway, proteoglycans in cancer, FoxO signaling pathway, etc. Molecular docking results suggested that sesamin was the most appropriate natural TNFis.

**Conclusion:** Our findings provide an important role and basis for further research on the molecular mechanism of SC treatment of RSA and drug development of TNFis.

## 1. Introduction

Recurrent spontaneous abortion (RSA), is a common pregnancy complication,which is defined as three or more clinically confirmed pregnancy failures, occurring in 12% of all pregnant women^[1]^, placing a serious burden on families and society.A wide array of studies confirms that RSA significantly affects the quality of life of women and their partners, increasing anxiety and depression in women with recurrent abortion^[2, 3]^. In addition, RSA is a risk factor for cardiovascular disease later in life^[4, 5]^.

The causes of recurrent abortion are complex, and the treatment options are mainly clinical psychological support and empiric therapy, including antithrombotic therapy and anti-immunotherapy, using aspirin, heparin, intravenous immunoglobulin (IVIG), together with other therapies, such as glucocorticoids, intra-lipid infusion, hormone supplementation with progesterone or dydrogesterone, granulocyte colony stimulating factor, and lymphocyte immunotherapy. However, the efficacy of these drugs is controversial, also having many adverse reactions including local allergy, liver damage, renal insufficiency, thrombocytopenia, osteoporosis, fatigue, lethargy, which confuse RSA patients^[6, 7]^. Therefore, it is essential to investigate effective and natural products to reduce pregnancy losses in patients with RSA.

In recent years, traditional Chinese medicine has become a research hotpots, as a popular western medicine supplement treatment of recurrent spontaneous abortion patients, with satisfactory curative effect, high safety and reliability. Cuscuta (Chinese name: Tusizi), the dried seeds of southern and Chinese dodder, have been used to cure abortion for many years ^[8]^. *Semen Cuscutae*, which is used as the core components of many classical herbal formula, has been reported to have functions of nourish the kidney and liver, anti-inflammatory properties, antibacterial, antioxidant, and antiulcer activities ^[9-10]^.

It was shown that lignans and flavonoids of Cuscuta sinensis extract could inhibit the release of NO and inflammatory factors PGE2, TNF-α and IL-6, thus suppressing the inflammatory response, and lignans could inhibit the expression level of cellular i NOS genes^[11,12]^. Although *SC* is widely used for the treatment of threaten abortion and RSA, the exact mechanisms still need to be further elucidated.

Tumor necrosis factor -α (TNF-α) plays an important role in cell proliferation, differentiation, apoptosis, immune regulation, and inflammatory induction,it has certain lethality to tumor cells, trophoblasts and embryonic tissues^[13, 14]^.Some researches have confirmed that serum level of TNF-a is higher at patients with recurrent abortion, suggesting that increased TNF-a may lead to abortion^[15]^.By down-regulating TNF-α to promote IL-6 expression,extract of SC can reduce maternal and fetal immune tolerance and maintain pregnancy continuation^[13]^. Recently, TNF-a has attracted more attention because of its important role in the inflammatory immune process, and inhibiting its biological activity can be used as a new direction for drug development.

Network pharmacology is an effective method for drug discovery. This paper aims to analyze the active components of *SC* by network pharmacology and screen the most stable compounds with TNF-a by molecular docking, to explore the mechanism of *SC* treatment of RSA and provide theoretical basis for drug development.

## 2. Materials and Methods

### 2.1 Database website

The databases used in this paper are shown in Table 1

**Table 1.**
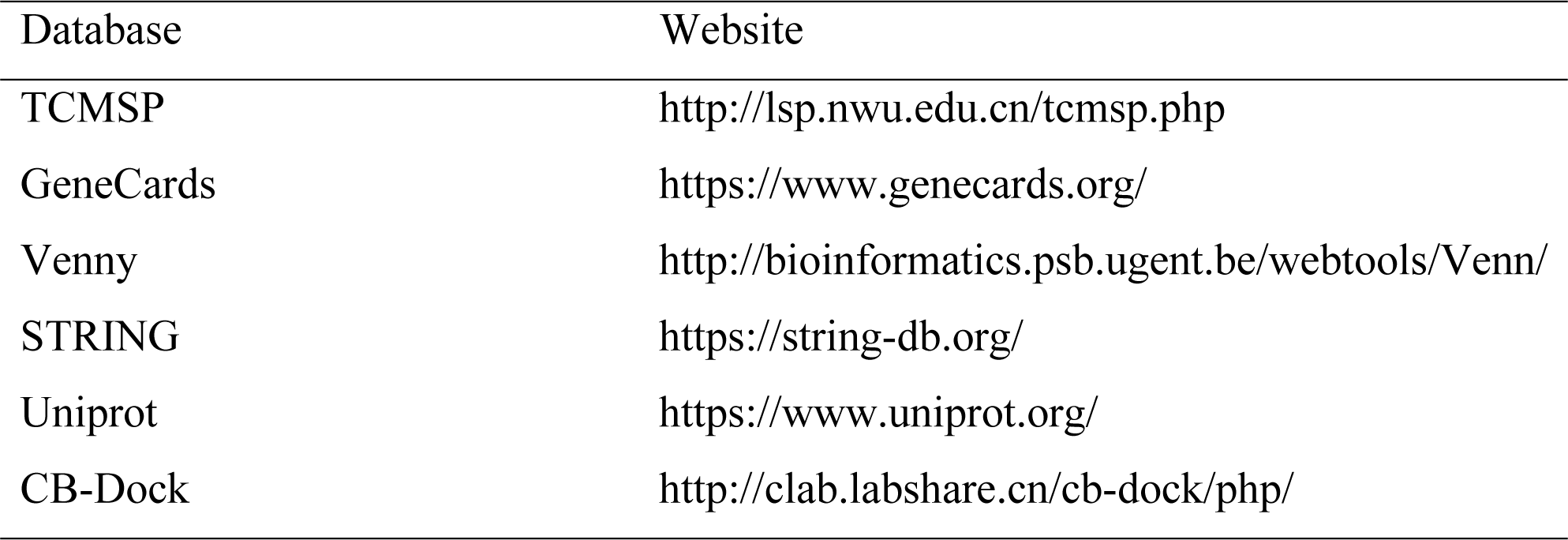
Database Website about this study.

| Database | Website |
| --- | --- |
| TCMSP | http://lsp.nwu.edu.cn/tcmsp.php |
| GeneCards | https://www.genecards.org/ |
| Venny | http://bioinformatics.psb.ugent.be/webtools/Venn/ |
| STRING | https://string-db.org/ |
| Uniprot | https://www.uniprot.org/ |
| CB-Dock | http://clab.labshare.cn/cb-dock/php/ |

### 2.2 Prediction of *SC* associated target genes and their intersection on RSA

The components of SC were obtained from the Traditional Chinese Medicine Systems Pharmacology Database and Analysis Platform (TCMSP, https://tcmspw.com/index.php) TCMSP is a Chinese herbal medicine system data platform on the drug-disease-target relationship, involving the pharmacokinetic properties of natural compounds, including oral bioavailability, intestinal epithelial permeability, drug similarity, blood-brain barrier, water solubility, etc ^[16]^.

According to literature reports and pharmacokinetic parameters, drug absorption, distribution, metabolism and excretion (ADME) are important factors affecting drug bio-activity. According to the TCMSP recommendations, compounds with oral bioavailability (OB) of 30% are well absorbed and metabolized slowly after oral administration. A drug similarity (DL) of 0.18 are chemically suitable for drug development^[17]^. Therefor,according to the relevant screening criteria (OB≥30%, DL ≥0.18), a total of 11 active ingredients and their target proteins were obtained from the TCMSP database. RSA related target genes were obtained from GeneCards. The combination target genes of *SC* treatment of RSA were obtained by Venny.

### 2.3 Construction of PPI network and functional enrichment analysis of related genes

STRING is a database for protein-protein interaction construction^[18]^. The combination target genes were inputted to the STRING database for further analysis analysis. The screening criteria was: Homo sapiens and confidence score greater than 0.4. The *SC*-compound-target-RSA network was visualization through Cytoscape software (version 3.6.1)^[19]^. The R software “cluster Profiler” and “pathview” packages were used to perform GO and KEGG pathway analysis and visualization.

### 2.4 Molecular docking technology

The active components selected from *SC* were ligated to TNF-a receptors by CB-Dock. CB-Dock is an online server that can automatically identify binding sites without predictive information, and then dock with AutoDock Vina according to the query results, with a success rate of about 70%, which is better than other commonly used blind docking tools^[20].^ PDB files for the TNF -α(protein ID: 2az5) was searched in the Uniprot database, while the active ingredients were download from TCMSP. They were posted on CB-Dock. The vina scores were used to evaluate the binding activity. The lesser the vina score, the more steady it is.

## 3. Results

### 3.1 Analysis of Cuscutae Semen compounds

According to the screening criteria, a total of 11 compounds were obtained from dodder. The specific information is shown in Table 2.

**Table 2.** Bioactive compounds of *Cuscutae Semen*.

| No | Molecule ID | Molecule name | Molecular weight | OB (%) | DL |
| --- | --- | --- | --- | --- | --- |
| 1 | MOL001558 | <u>sesamin</u> | 354.38 | 56.55 | 0.83 |
| 2 | MOL000184 | <u>NSC63551</u> | 412.77 | 39.25 | 0.76 |
| 3 | MOL000354 | <u>isorhamnetin</u> | 316.28 | 49.60 | 0.31 |
| 4 | MOL000358 | <u>beta-sitosterol</u> | 414.79 | 36.91 | 0.75 |
| 5 | MOL000422 | <u>kaempferol</u> | 286.25 | 41.88 | 0.24 |
| 6 | MOL005043 | <u>campest-5-en-3beta-ol</u> | 400.76 | 37.58 | 0.71 |
| 7 | MOL005440 | <u>Isofucosterol</u> | 412.77 | 43.78 | 0.76 |
| 8 | MOL005944 | <u>matrine</u> | 248.41 | 63.77 | 0.25 |
| 9 | MOL006649 | <u>sophranol</u> | 264.41 | 55.42 | 0.28 |
| 10 | MOL000953 | <u>CLR</u> | 386.73 | 37.87 | 0.68 |
| 11 | MOL000098 | <u>quercetin</u> | 302.25 | 46.43 | 0.28 |

### 3.2. Drug -Disease-Compounds-Target gene network

128 target genes corresponding to 10 active components of SC were obtained from TCMSSP database. (sophranol had no direct targets).Similarly,there were 1986 target genes of RSA obtained by searching GeneCard database excluding duplicate genes. 79 potential target genes were obtained by crossing RSA and SC target genes by Venny software(Fig 1). The PPI network map was obtained by importing 79 target genes into STRING database, which was visualized using Cytoscape software(Fig 2 A), the top 10 targets were found by further analysis, namely TNF, IL-6, TP53, IL1B, EGFR, JUN, CASP3, ESR1, PPARG, PTGS2.(Fig 2 B). The above results were input into the Cytoscape to obtain a *SC*–RSA-compounds-gene network (Fig 3).

**Fig 1.**
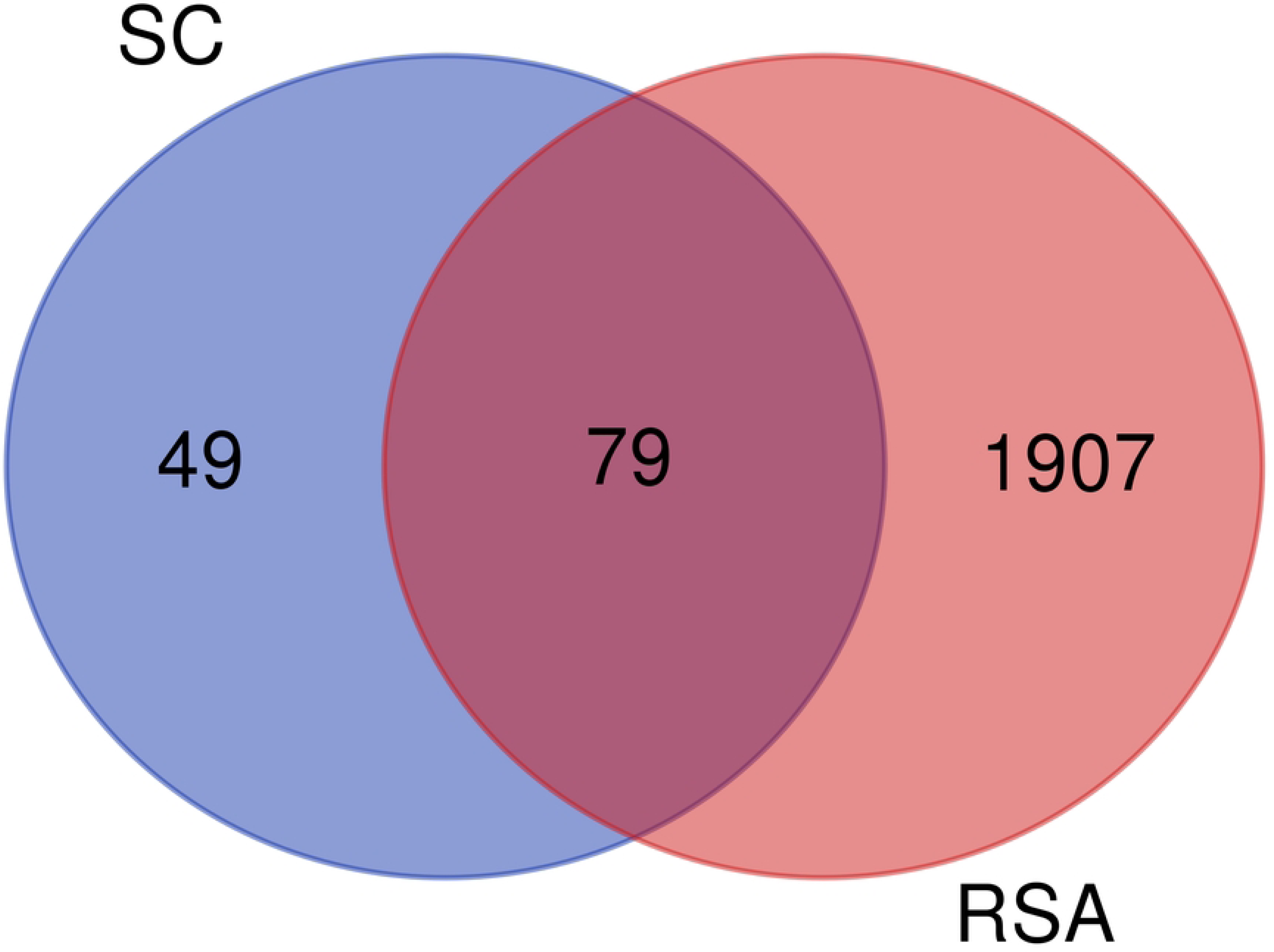
Venn diagram of 79 target genes: potential targets for SC treatment of RSA.

**Fig 2.**
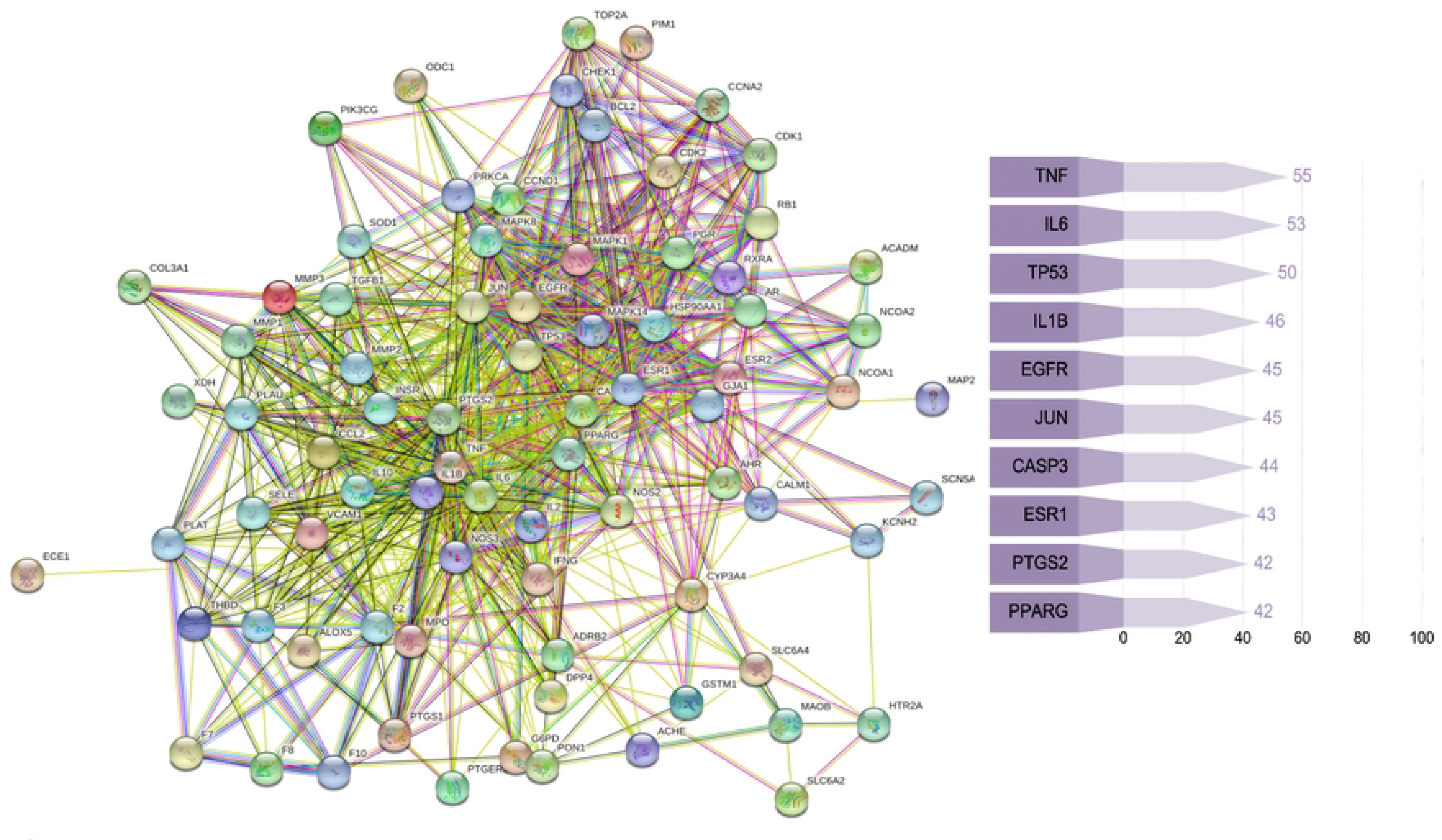
Potential target genes and PPI network map of SC therapy for RSA(A:The PPI network map of 79 target genes B:list of the top 10 genes of PPI network map)

**Fig 3.**
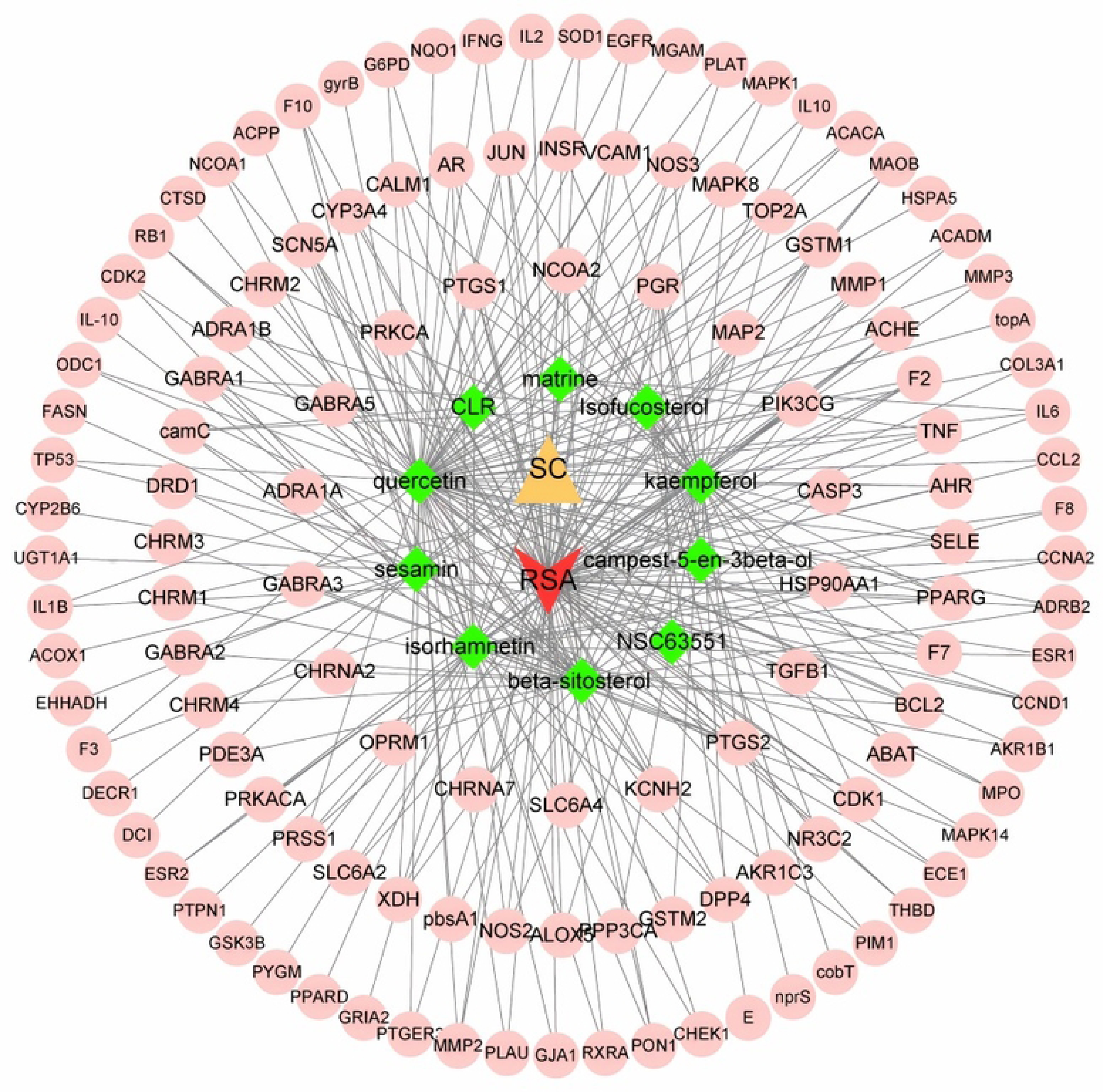
S*C*–RSA-compounds-potential target gene network map. Yellow triangle represents SC, red v represents RSA, green diamonds mean the SC’s active ingredients, while the pink circles mean 79 overlapping target genes.

### 3.3 Enrichment analysis of GO and KEGG pathway

57 biological function terms were obtained by GO enrichment analysis, and the threshold value was P<0.05. The top 20 catalogs were chosen for scatter plots (Fig 4). The biological functions of these genes are mainly positive regulation of RNA polymerase II promoter transcription, positive regulation in transcription, DNA templating, oxidation-reduction process, positive regulation in gene expression, inflammatory response.

**Fig 4.**
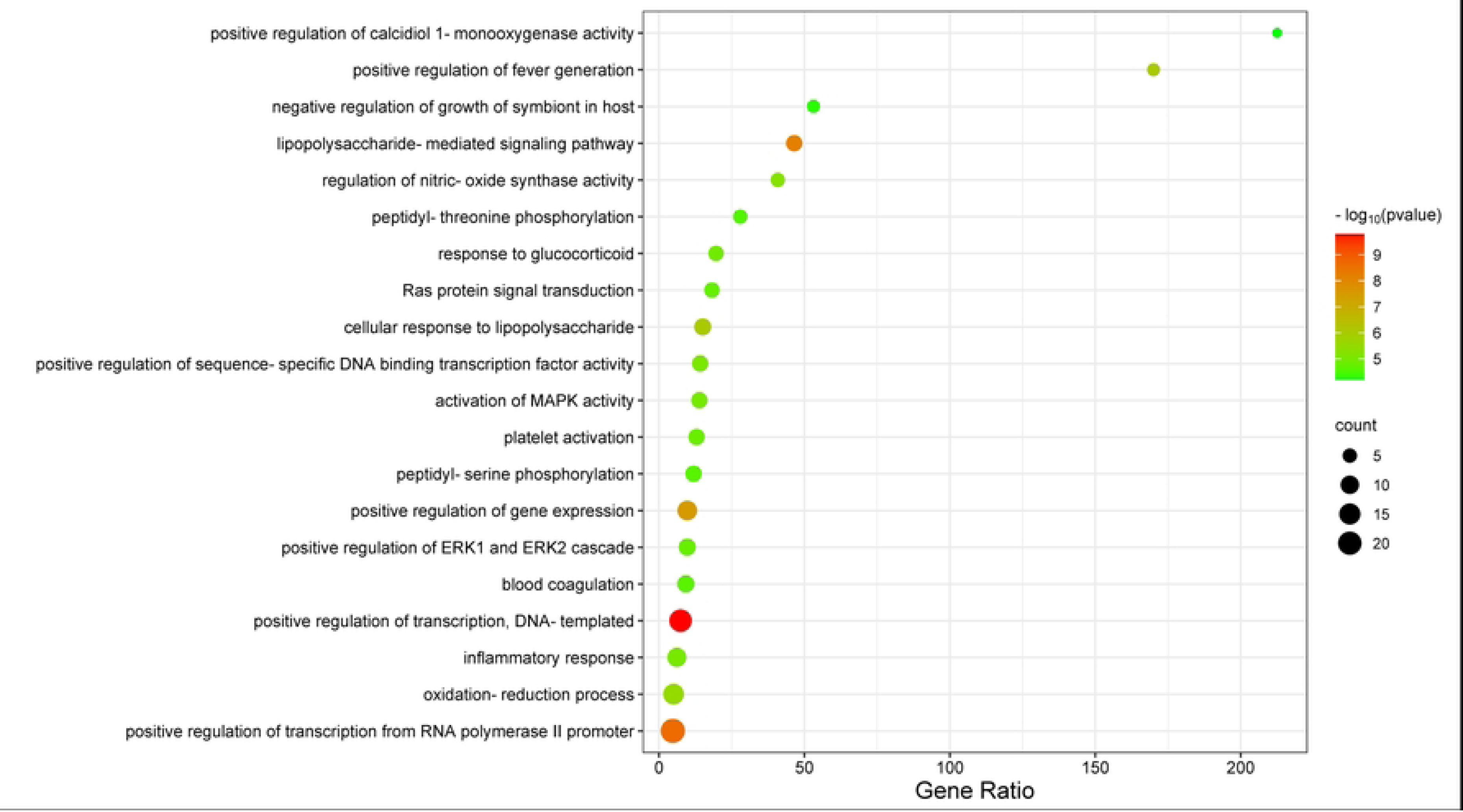
The top 20 biological functions.

To further understand the mechanism of *SC* in RSA treatment, 82 signaling pathways were obtained by enrichment analysis of KEGG pathway (p < 0.05). The scatterplot showed the first 15 essential signal pathways (Fig 5), part of which are strongly related to RSA, for instance TNF signaling pathway, HIF-1 signaling pathway, Estrogen signaling pathway, FoxO signaling pathway, NOD-like receptor signaling pathway. Furthermore, signaling pathway of vital TNF-α is shown in Fig 6.

**Fig 5.**
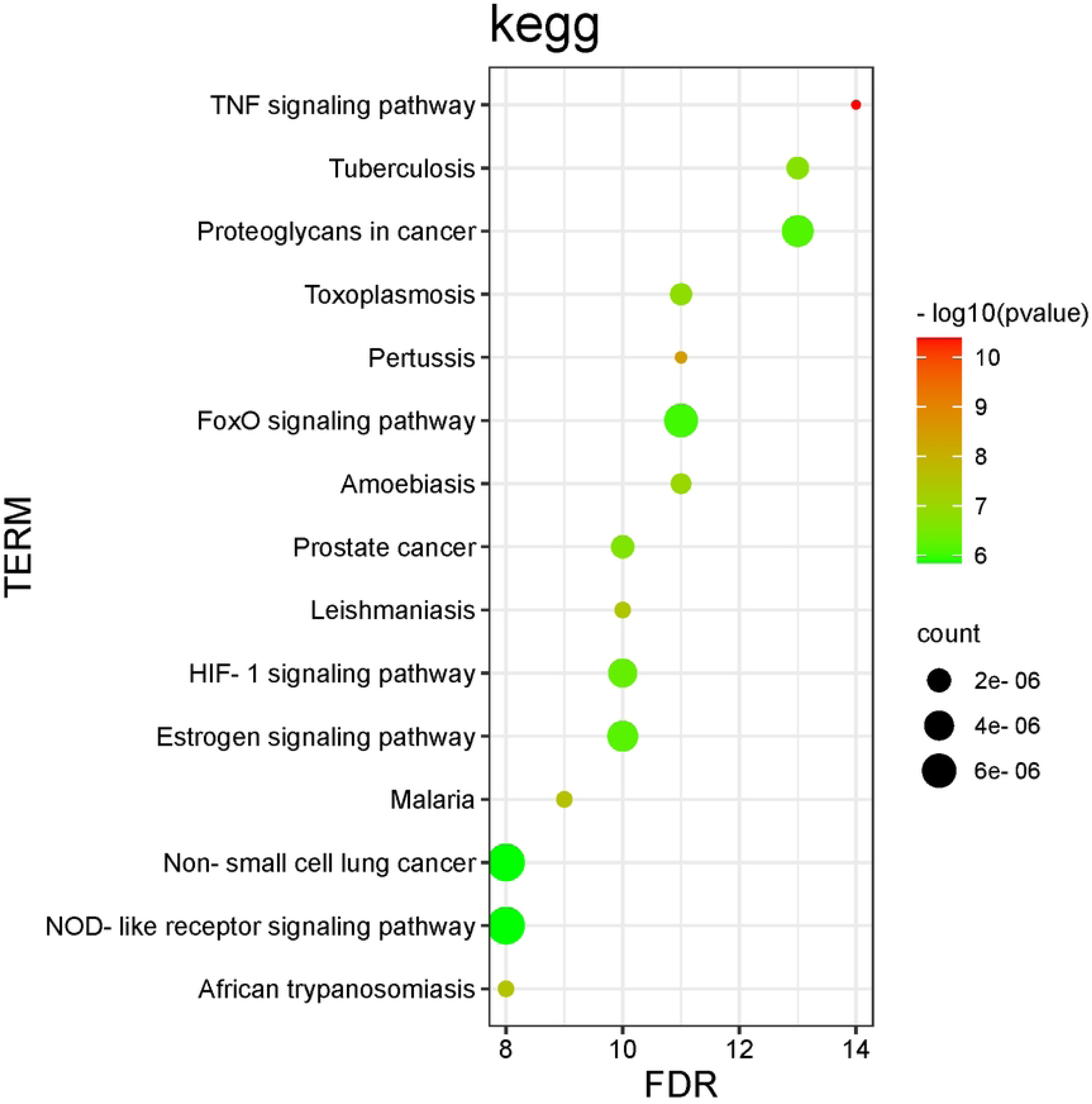
The top 15 signal pathways for KEGG enrichment of core targets.

**Fig 6.**
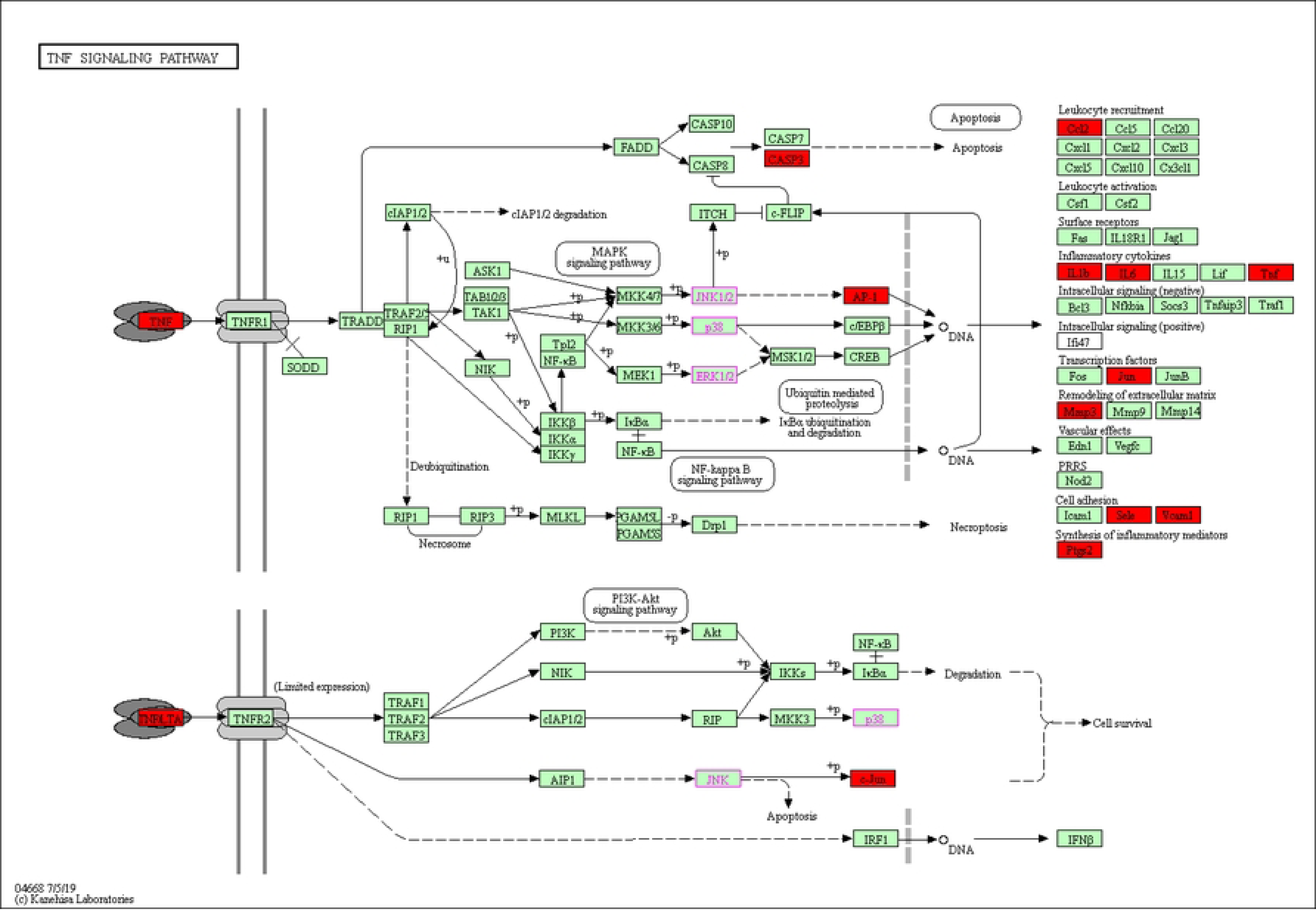
Core target TNF signaling pathway. Red color represents SC’s action targets in the network.

### 3.4 Binding force between TNF-α and the active constituent

10 active ingredients were found to bind to TNF-α in various degrees **(Table 3).** Venous score signify binding capacity, with lower scores, the compound and acceptor will react to each other more strongly and steadily. Sesamin had the highest Venous scores with TNF-a,the results demonstrated that sesamin might have the strongest and consistent affinity for TNF-a among the ten compounds assessed.The molecular binding of the compounds to TNF-α is demonstrated in Fig 7.

**Fig 7.**
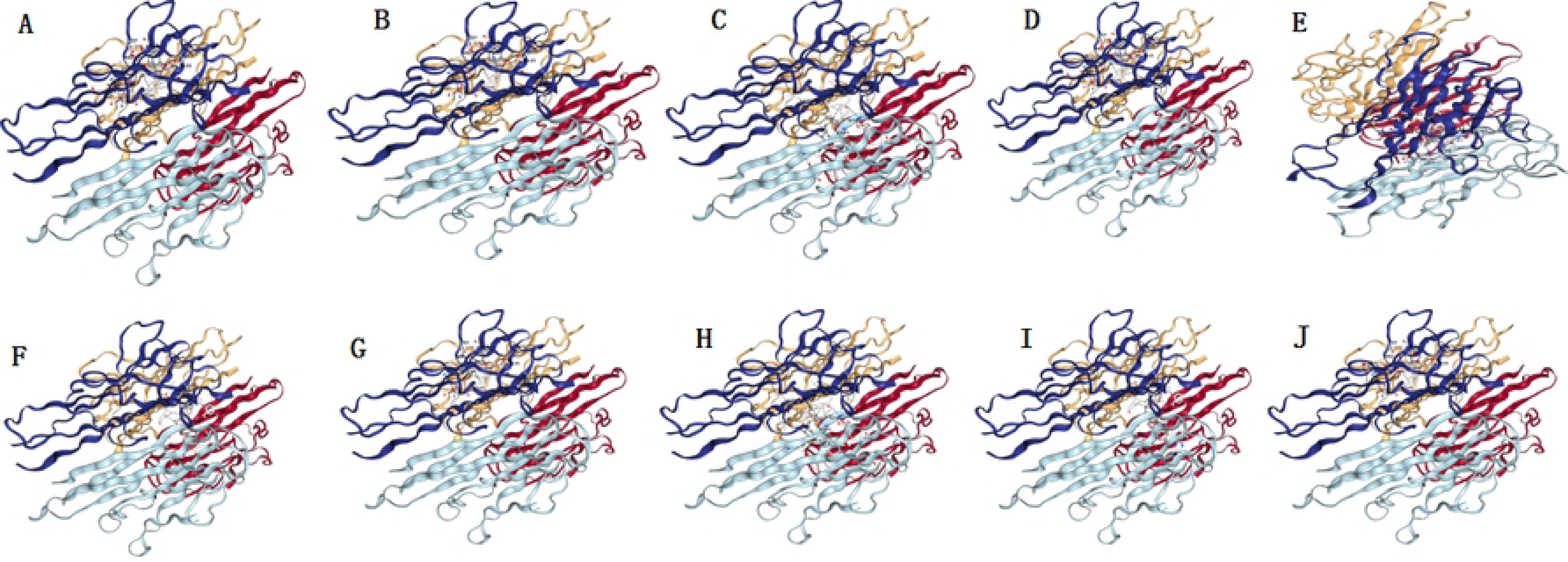
The three dimensional map of the binding of compounds. (A) quercetin, (B)kaempferol, (C) beta-sitosterol, (D)isorhamnetin, (E)NSC63551, (F) CLR,(G) matrineand, (H)Isofucosterol,(I)campest-5-en-3beta-ol and (J)sesamin with TNF-α.

**Table 3.**
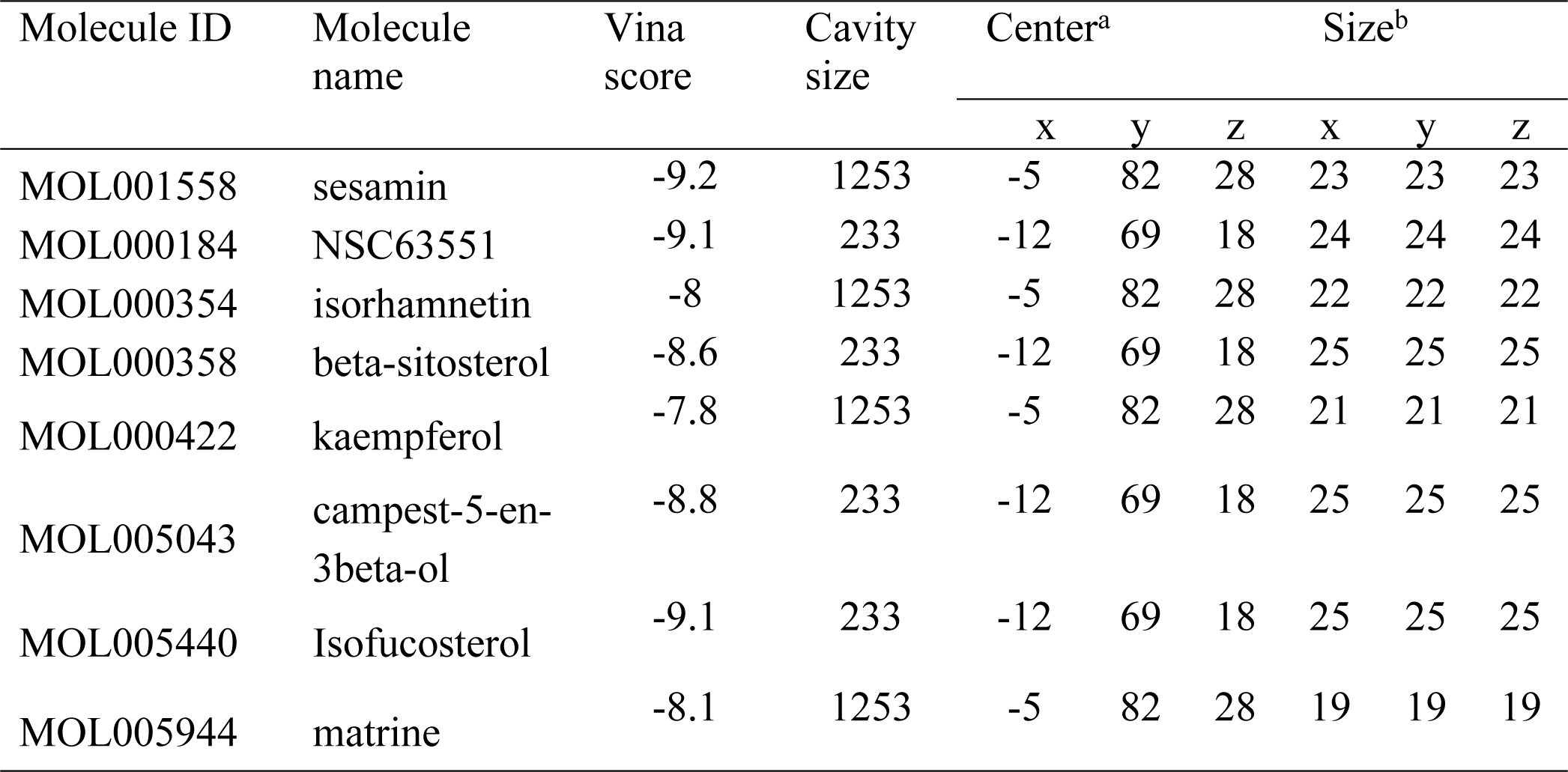
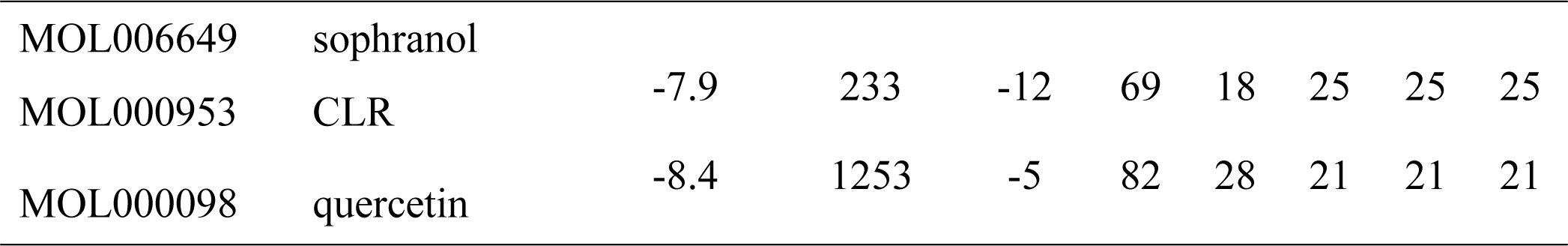
Molecular docking parameters and results of the binding of SC active ingredients to TNF-α.

## 4.Discussion

RSA affects 1-2% of couples and causes great psychological and spiritual trauma to women, as well as financial burden. The incidence of RSA is difficult to estimate due to differences in definitions and criteria. The causes of RSA are complex and numerous. The known causes include female age, genetic,anatomical and chromosomal abnormalities,endocrinology, infection, placental anomalies, smoking and alcohol consumption, psychological factor and so on^[21]^. In form of herbal mixtures, Cuscuta has been known as a cure to many diseases, for instance, oligospermia ^[22]^, osteoporosis ^[23]^, premature ovarian failure ^[24]^, Induction of sleep ^[25]^, habitual abortions ^[26]^, and neurasthenia^[27]^.

In this study, 10 active ingredients (including flavonoids, glycosides, alkaloid, and steroids) from *SC* play an important role in the therapy of RSA. Hyperoside is a flavonoid, which belongs to polyphenolic compounds. Some studies have reported that hyperoside always show a protective response on pregnancy loss in rats by enhancing autophagy and inhibiting inflammation. This effect may be related to the inhibition of mTOR/S6K^[28]^.Aberrant dendritic cell (DC) activities and differentiation have been identified to be associated with RSA. It has been reported that baicalin attenuates embryonic resorption in RSA mice by inhibiting STAT5-ID2 expression and reversing the expression of traditional DCs to plasmacytoid DCs and functional molecules^[29]^.There is increasing evidence that T-helper cell imbalance is related to RSA. One study suggested that CD4 T-helper cells and circulating/uterine natural killer (NK) cell populations are associated with RM and implantation failure^[30]^. Th1/Th2 imbalance can lead to recurrent miscarriage^[31]^, as confirmed by Niu Y, matrine can regulate the balance through NF-κ B signaling^[32]^. Some studies have also confirmed that matrine can regulate Th1/Th2 balance through upregulated IFN-γ and downregulated TGF-βexpression^[33]^.Overall,these compounds are related to multiple proteins and signal pathways, which are of significant potential research value on RSA and deserve further investigation. Moreover, as shown in Figure 2, results of PPI indicated that 128 target proteins are not independent of each other, but are linked and interact.TNF, IL-6, TP53, IL1B, EGFR, JUN, CASP3, ESR1, PPARG, PTGS2 were top 10 hub genes.As seen in Figure3, many compounds can be involved in RSA by regulating multiple target genes,such as TGFβ1, PTGS2, KCNH2, OPRM1. Enrichment analysis of GO and KEGG showed that 57 biological functions and 82 signal pathways took part in the development of RSA, which may be the mechanism of treatment of recurrent abortion by *SC*.The levels of a variety of cytokines in RSA patients are changed, such as TNF-a and interleukin family, which may be involved in the occurrence of RSA,^[34]^. Studies have confirmed that IL-4, IL-10 promotes embryonic development and placentation, while IFN-c and TNF-a inhibit trophoblast growth and differentiation, having the opposite effect^[35]^.The FOXO signaling pathway regulates a variety physiological events, for example, cell-cycle control,apoptosis, oxidative stress resistance and glucose metabolism. There are no reports on the relationship between FOXO signaling pathway and RSA. FOXO signaling pathway may have some relationship with SA in the above aspects, but further studies are needed.

TNF signaling pathway is a classical signaling pathway involvedin the occurrence and development of many diseases. Expression of inhibiting TNF-a and the symptoms of anti-TNF-a antibody therapy that alleviate inflammatory diseases^[36]^.The active components of *SC* can regulate sex hormone abnormalities, and reduce oxidative stress, ER stress and cell apoptosis by lower the TNF-a’s level and IL-6^[37]^.TNF-a is significant in immunity and inflammation at the maternal-fetal interface. Recently, the relationship between the TNF-α and RSA has received much attention.Further studies have found that inhibiting the Th1’s expression cytokines, for instance, TNF-a is beneficial to create a microenvironment at the feto-maternal interface and maintain pregnancy^[38]^. At present, anti-TNF-α drugs, such as etanercept (ETA), infliximab (IFX),adalimumab (ADA), golimumab and certolizumab have been widely used in clinical inflammatory diseases,but, these drugs are associated with adverse pregnancy outcomes. For example, IUGR, SA, and preterm birth are the most common adverse pregnancy outcomes associated with anti-TNF-a drugs^[39-42]^.In addition, severe infection of pregnant women, gestational diabetes mellitus, stillbirth, premature rupture of membranes and hypertension during pregnancy have also been reported in some reports^[43-45]^. At present, there is a lack of expert consensus and guidelines on the use of these drugs during pregnancy.

In our work, sesamin with good OB and DL properties was selected from *SC*. Moreover, it had strongest binding ability to TNF-α molecule. Sesamin originates from the western regions of ancient China and belongs to the flax seed family^[46]^. Sesamin can significantly reverse intestinal ischemia-reperfusion -increased IL-6,TNF-α and IL-1β in serum, exerts anti-inflammatory and antioxidant effects through Nrf2/HO-1/NQO1 signaling pathway^[47]^. In vitro model studies suggest that sesamin has a protective effect on cartilage destruction induced by the combination of tumor necrosis factor-α and ondomasin-M^[48]^. These results suggest that sesamin can inhibit TNF-α and may be a promising alternative treatment for patients with RSA.Further studies need to be completed for experimental validation.

Taking these findings into consideration provides a theoretical basis for further development of natural TNFis and the development of new anti-inflflammatory immune drugs using SC as the base substance.However, this paper has some limitations: 1. The absorption of compounds in humans is not limited to OB; 2. Interactions between active ingredients are not taken into account

## Acknowledgments

The authors are grateful to all members of the Department of Gynecology and Obstetrics, the People’s Hospital of China Three Gorges University/the First People’s Hospital of Yichang.We are grateful to the public database for providing meaningful datasets for our research.

## Authors’ Contributions

Wenfei Zheng and Manshu Lei conceived and designed the research, and wrote the manuscript. Yao Yao, Jingqiong Zhan applied the software. Yiming Zhang, Feng Huang completed visualization. Wenfei Zheng supervised the study and reviewed the manuscript. All authors have read and approved the final manuscript.

